# Heterogeneous adaptation of cysteine reactivity to a covalent oncometabolite

**DOI:** 10.1101/2020.06.25.171843

**Authors:** Minervo Perez, Daniel W. Bak, Sarah E. Bergholtz, Daniel R. Crooks, Youfeng Yang, Eranthie Weerapana, W. Marston Linehan, Jordan L. Meier

## Abstract

Metabolism and signaling intersect in the genetic cancer syndrome hereditary leiomyomatosis and renal cell carcinoma (HLRCC), a disease in which mutation of the TCA cycle enzyme fumarate hydratase (FH) causes hyperaccumulation of fumarate. This electrophilic oncometabolite can alter gene activity at the level of transcription, via reversible inhibition of epigenetic dioxygenases, as well as posttranslationally, via covalent modification of cysteine residues. To better understand how metabolites function as covalent signals, here we report a chemoproteomic analysis of a kidney-derived HLRCC cell line. Building on previous studies, we applied a general reactivity probe to compile a dataset of cysteine residues sensitive to rescue of cellular FH activity. This revealed a broad upregulation of cysteine reactivity upon FH rescue, caused by an approximately equal proportion of transcriptional and posttranslational regulation in the rescue cell line. Gene ontology analysis highlights new targets and pathways potentially modulated by FH mutation. Comparison of the new dataset to literature studies highlights considerable heterogeneity in the adaptive response of cysteine-containing proteins in different models of HLRCC. Our analysis provides a resource for understanding the proteomic adaptation to fumarate accumulation, and a foundation for future efforts to exploit this knowledge for cancer therapy.

## Introduction

Metabolites play diverse roles in cellular homeostasis, acting as transcription factor ligands, secondary messengers, feedback inhibitors, and allosteric effectors of enzyme function.^1–3^ An emerging function by which metabolites modulate cell signaling is via covalent protein modification due to their intrinsic reactivity.^4–5^ For example, in the genetic cancer predisposition syndrome hereditary leiomyomatosis and renal cell carcinoma (HLRCC) mutation of the tricarboxylic acid (TCA) cycle enzyme fumarate hydratase (*FH*) causes the hyperaccumulation of fumarate, an electrophilic “oncometabolite.” Reaction of cysteines with fumarate’s α,β-unsaturated double bond results in protein S-succination, a non-enzymatic posttranslational modification (PTM) that can substantially modulate protein function.^6–8^ In addition to direct modification, fumarate accumulation can also indirectly alter cysteine activity, though redox stress that causes oxidative cysteine modifications,^9–11^ or transcriptional changes that upregulate or downregulate the expression of cysteine-containing proteins (Fig. 1a).^12–16^

**Fig. 1.**
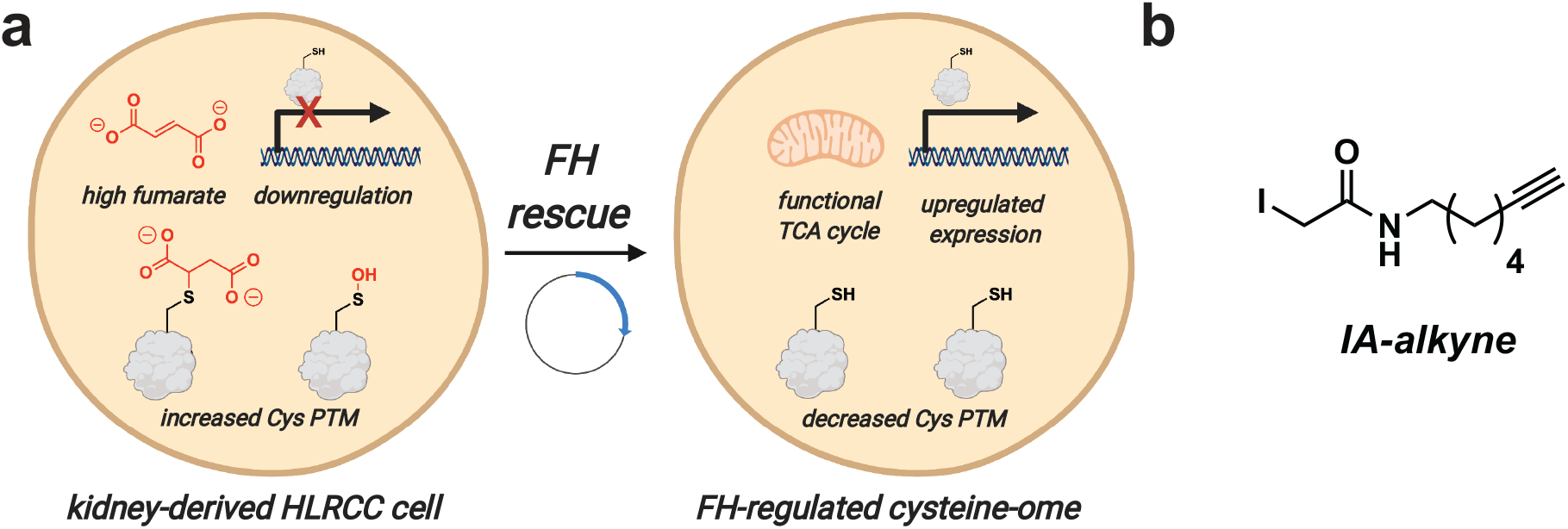
(a) Left: Inactivation of FH in the genetic cancer syndrome HLRCC causes high levels of fumarate to accumulate, which can decrease cysteine reactivity by downregulating protein expression and increasing protein posttranslational modification. Right: FH rescue can reverse many of these changes, which can be identified based on their altered cysteine reactivity. (b) IA-alkyne, an enrichable reactive electrophile used to quantify cysteine reactivity in this study.

## Results

To better understand fumarate’s potential as a metabolic signal our lab recently reported a quantitative chemoproteomic approach to study conditional cysteine reactivity in HLRCC.^17^ This approach uses a chemical probe, iodoacetamide alkyne (IA-alkyne), to substoichiometrically label cysteines across the proteome in isogenic FH-deficient (*FH*−/−) and wild-type (*FH*+/+) rescue cell lines (Fig. 1b).^18^ Conjugation to azide-biotin tags using click chemistry followed by enrichment, cleavage, and quantitative LC-MS/MS analysis enables comparison of how the reactivity of individual cysteine residues are regulated by FH rescue and subsequent reduction in intracellular levels of fumarate. In our initial study we applied this method to quantify *FH*-regulated cysteine residues in an immortalized cell line derived from an HLRCC metastasis. This led to the identification of new candiate targets of S-succination, as well as evidence that protonation of fumarate is necessary for it to act as a covalent modifier in HLRCC.^17^

Indirect readouts such as HIF accumulation,^19^ oxidative respiration,^20^ and anti-S-succination immunoblotting^17^ indicate that fumarate accumulates at different levels in cell lines derived from primary HLRCC tumors as compared to distal metastases. This suggests the *FH*-regulated cysteine-ome of different tumors may display substantial heterogeneity, potentially contributing to distinct biology, diagnostic markers, or therapeutic vulnerabilities. To investigate this hypothesis, we applied our chemoproteomic approach to an immortalized cell line derived from the kidney of an HLRCC patient.^20^ Briefly, a kidney-derived HLRCC cell line (UOK268, *FH−/−*) and a rescue line in which the FH gene had been re-introduced (UOK268, *FH+/+*) were cultured on large-scale and proteomes were isolated from ~30-40 million cells by mild sonication. Paired samples were IA-alkyne labeled, conjugated to a UV-cleavable biotin azide via click chemistry, enriched, isotopically tagged, pooled, and analyzed by LC-MS/MS (Fig. 2a). Analysis of the intensity ratio of MS1 spectra of light/heavy (L/H) isotopic pairs in IA-alkyne treated samples was used to determine relative cysteine reactivity between the two samples. An R value of 1 indicates a cysteine was equally reactive in *FH−/−* and *FH+/+* samples, whereas an increased R value of 2 indicates a cysteine’s reactivity was recovered by 50% upon *FH* rescue (based on the formula “relative reactivity (%) = [1-(1/R)]*100%”; Table S1).

**Fig. 2.**
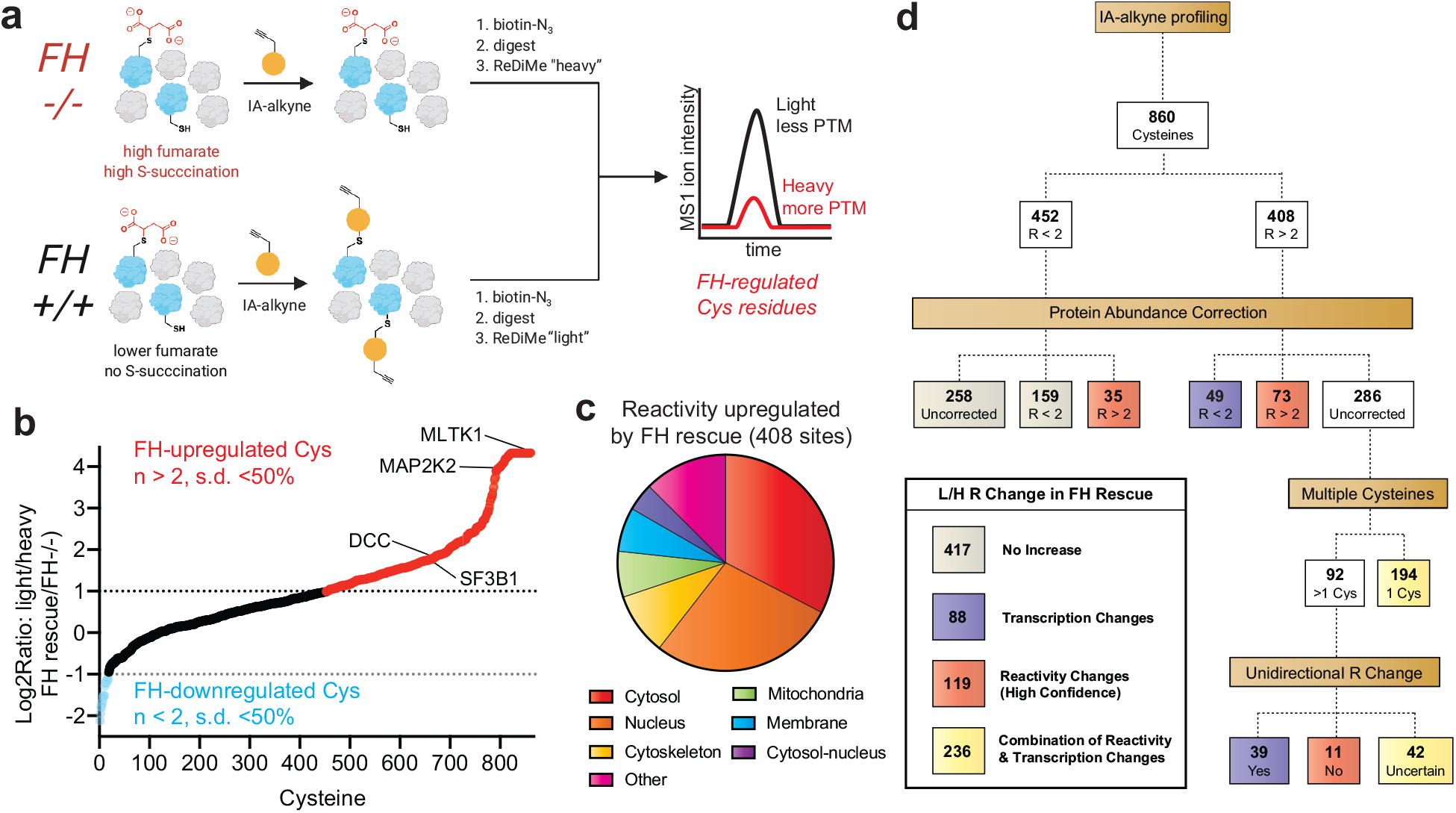
Defining the consequences of FH-rescue in HLRCC via quantitative cysteine reactivity profiling. (a) General workflow for reactive cysteine profiling experiment. FH−/− = FH mutant cells, FH+/+ = FH rescue cells, IA-alkyne = iodoacetamide alkyne. (b) Graph of quantified cysteine reactivity in FH−/−(heavy) and FH+/+ (light) UOK268 cell lines, sorted from high to low peptide R values. (c) Cellular localization of cysteine residues upregulated by FH rescue (R>2, SD<50%). (d) Flow chart describing how protein abundance correction and the presence of multiple cysteine residues in a single protein were used to define cysteine reactivity changes likely to be driven by FH-dependent transcription (purple), FH-dependent reactivity (red), and a combination thereof (yellow).

Performing four independent replicate measurements of cysteine reactivity in paired *FH+/+* and *FH−/−* kidney-derived UOK268 cell lines enabled the analysis of cysteine reactivity across 860 individual residues with high confidence (n >2, standard deviation of R (reactivity) of < 50%) (Fig. 2b, Table S2). Assessing the directionality of cysteine reactivity changes between the isogenic cell lines, we found the reactivity of 408 cysteines were up-regulated ≥2-fold upon FH rescue (Fig. 2b, red), while the reactivity of only 17 cysteines were down-regulated to this extent (Fig. 2b, blue). This is consistent with our expectation of increased cysteine reactivity in *FH* rescue (*+/+)* due to decreased accumulation of the electrophilic oncometabolite fumarate.

Interestingly, the cysteines whose reactivity was most strongly increased upon FH rescue predominantly mapped to the nucleus and cytosol (Fig. 2c), while the relatively few cysteines whose reactivity was decreased by FH rescue were enriched in mitochondrial proteins (Fig. S1a). Hypothesizing that the latter effect may be due to compensatory upregulation of mitochondrial proteins in FH−/−cells, we applied two complementary approaches to examine the relationship between protein expression and cysteine reactivity in our dataset (Fig. 2d). First, we used reductive dimethyl (ReDiMe) labeling to quantitatively assess changes in overall protein abundance between the two cell lines (Table S3).^21^ We obtained quantitative abundance data for the parent proteins of one-third of cysteines whose relative reactivity was measured in UOK268 *FH−/−* and rescue cell lines (Table S4). Overall, relative cysteine reactivity and protein abundance showed a strong correlation in the two HLRCC cell lines (Pearson’s r = 0.57, Fig. S1b). Cysteines exhibiting the greatest reactivity increases upon *FH* rescue (R≥2) also showed the most augmented expression in this condition, consistent with the overall trend (Fig. S1c). Focusing on cysteine residues with increased reactivity upon FH-rescue (R≥2), we found 49 were also strongly upregulated (≥2-fold decreased) at the protein level (Fig. 2d, Table S4), suggesting upregulated protein expression. Conversely, the FH-dependent reactivities of 73 cysteines was relatively unchanged, while an additional 35 shifted from R≤2 to R≥2 when correcting for protein abundance, consistent with the potential for these residues to be targets of FH-dependent posttranslational modification (Fig. 2d, Table S4).

To extend this analysis to *FH*-regulated (R≥2) cysteines whose proteomic abundance was not sampled by ReDiMe we developed a second approach, examining cases where multiple IA-alkyne quantifiable cysteine residues were identified within a single polypeptide (Fig. 2d, Table S5). Our hypothesis was that changes in expression levels should cause a consistent change in reactivity across multiple quantified cysteine residues of a protein, while changes in cysteine post-translational modifications, such as S-succination or cysteine oxidation, may alter the reactivity of only a subset of sites.^22^ Applying this approach, we observed consistent (SD≤50%) unidirectional changes in cysteine reactivity for 39 of these *FH*-regulated (R≥2) cysteines and distinct reactivity changes (SD≥50%) for 11 additional cysteine residues, with the remainder (42) falling in between (Fig. 2d, Table S5). For example, C154, C380, and C410 of UQCRC1 display very similar R values (2.61, 2,07, and 2.73 respectively) despite residing in distinct domains of the protein. In contrast, two unique cysteine residues were quantified within the microsomal protein LRRC59, one of which displays distinct FH-dependent reactivity (C277, R=14.86; C48, R=1.95). These studies suggest cysteine reactivity changes sampled by IA-alkyne in this isogenic HLRCC model are caused by a combination of gene expression changes (88 cysteines) and FH-dependent posttranslational protein modifications (119 cysteines).

Our comprehensive chemoproteomic dataset affords a unique opportunity to define how cellular pathways respond to re-introduction of the FH tumor suppressor at both the level of protein expression and cysteine reactivity. Focusing first on protein expression (Figure 3, Table S3), gene ontology analysis using the informatic tool DAVID^23^ found *FH* rescue upregulated the expression of proteins involved in cell-cell adhesion (FLNA, MYH9, PFN1, KIF5B), protein translation (EIF4G2, RPL12, RPL13, RPS25, RPLP1), and glycolysis (ALDOA/C, ENO1, GPI, GAPDH, PFKP, PGK1, PGAM1, TPI1, LDHA/B), while downregulating mitochondrial enzymes involved in the TCA cycle and redox homeostasis (Fig. 3b-c). HLRCC tumors are well-known for their metastatic potential,^16, 24^ and the upregulation of cell adhesion proteins suggests *FH* rescue may reverse this phenotype. In contrast, the upregulation of glycolytic enzymes upon re-introduction of *FH* activity was unexpected. Previous studies have established that glycolytic gene expression is upregulated in HLRCC tumors and tumor-derived cells,^20^ and chemoproteomic profiling analysis also found that rescue of *FH* in the metastasis-derived UOK262 HLRCC cell line downregulated many glycolytic enzymes at the level of protein expression, suggesting that UOK262 cells may show a greater degree of metabolic flexibility.^17^ The glycolytic upregulation of UOK268 FH rescue cells highlights a potential caveat of our model, which is that re-introduction of FH into transformed cells may recapitulate only a subset of differences between HLRCC tumors and healthy tissues, due to irreversible signaling changes and genomic insults that accompany malignant transformation. Our results indicate that restoring the respiratory capability of kidney-derived UOK268 cells does not necessarily “cure” the heavy reliance of these cells on glycolytic enzymes. FH rescue disproportionately affected the levels of enzymes involved in oxidation-reduction processes, causing ≥2-fold downregulation of eight mitochondrial NAD(P)H producing enzymes: OGDH, MDH2, IDH3A (all members of the TCA cycle), ALDH2, GLUD1, FDXR, HIBADH, ME3 (Table S3, Fig. 3d-e), consistent with previous observations of FH-regulated changes in redox homeostasis.^9–10, 25^ Our studies, together with the literature, are consistent with a model in which FH inactivation causes a dependence on enzymes involved in glycolytic metabolism and the production of reducing equivalents in the mitochondria, only the latter of which is reversed by re-introduction of functional FH activity.

**Fig. 3.**
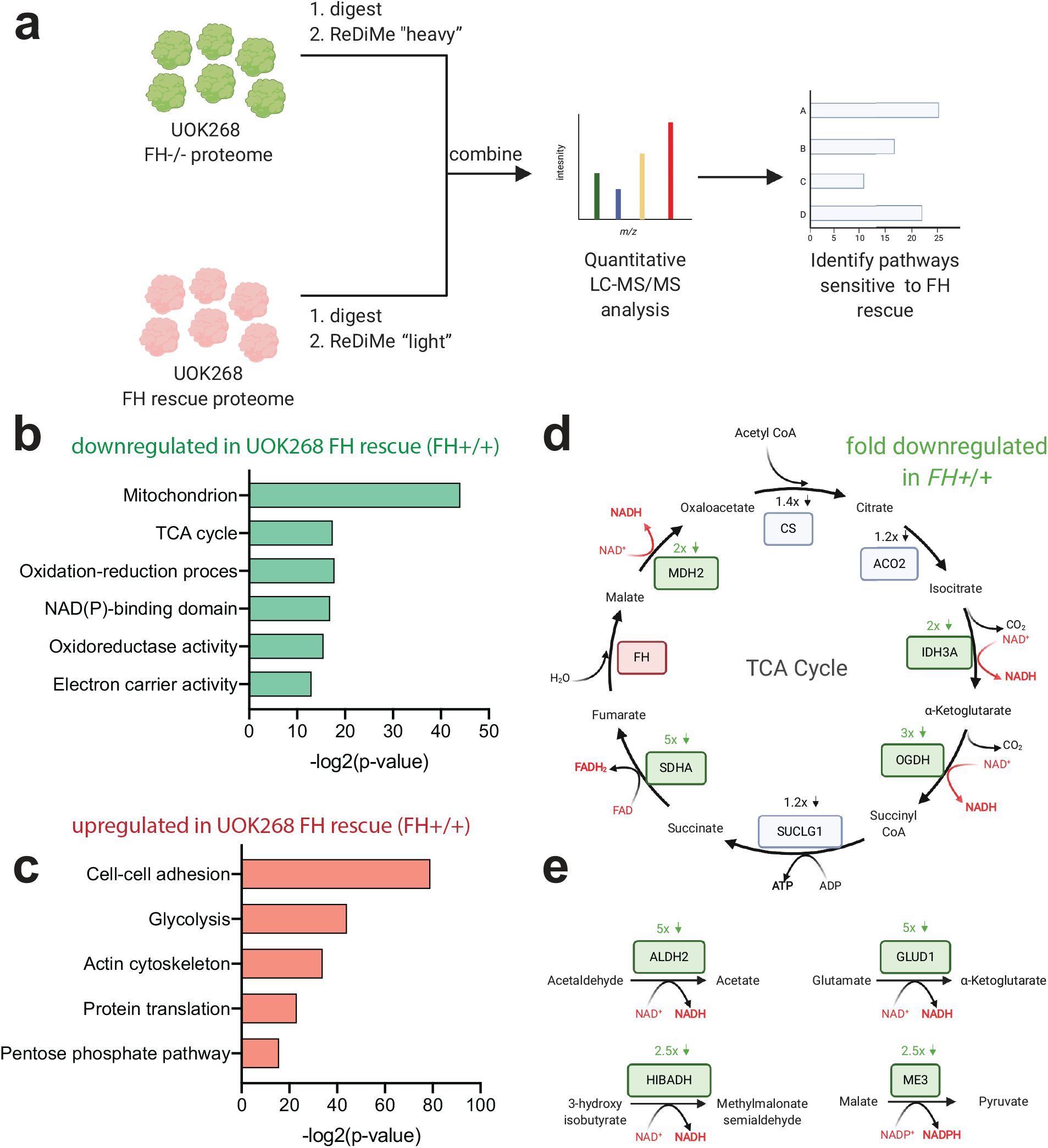
Defining the consequences of FH-rescue in HLRCC at the whole proteome level via quantitative reductive dimethyl (ReDiMe) labeling. (a) Workflow for whole proteome ReDiMe analysis of UOK268 HLRCC cells. (b) Gene ontology terms encompassing proteins found to be downregulated upon FH rescue in kidney-derived HLRCC cells. (c) Gene ontology terms encompassing proteins found to be upregulated upon FH rescue in kidney-derived HLRCC cells. (d) Influence of FH rescue on TCA cycle enzyme expression. Genes boxed in green are downregulated >2-fold in the FH rescue (FH+/+) UOK268 cell line relative to mutant (FH−/−). (e) Influence of FH rescue on NAD(P)H producing mitochondrial metabolic enzymes. Genes boxed in green are downregulated >2-fold in the FH rescue (FH+/+) UOK268 cell line relative to mutant (FH−/−).

Next, we explored changes in cysteine reactivity not explained by altered protein abundance between the mutant and rescue UOK268 cell lines, which represent candidate sites of FH-dependent posttranslational modification (Fig. 4a) (Tables S4-6; Fig. 4a). Here, we focused on 119 “high confidence” residues which displayed an R≥2 following correction for protein abundance or multiple cysteines (Fig. 2d, red), as well as an additional 236 “lower confidence” cysteines whose parent proteins were not quantified by whole proteome LC-MS/MS analysis, and whose reactivity changes thus may be driven by <u>either</u> altered expression or posttranslational modifications (Fig. 2d, yellow; Tables S4-6). To identify functional cysteine modifications with potential cancer-relevance, we employed a two-step analysis. First, the parent proteins of these 355 candidate FH-regulated cysteine residues were queried for evidence of involvement in human cancer using the Network of Cancer Genes database.^27–28^ Hits from this analysis were then assessed for conservation of the candidate modified cysteine using the online informatic tool Mutation Assessor.^26^ Applying this strategy we identified 55 genes known to be deleted, overexpressed, or amplified in human cancer that harbored a total of 69 candidate FH-regulated cysteine residues. In 8 instances, mutation of the FH-regulated cysteine was predicted to highly perturb the function of the parent protein, while mutation of a further 23 cysteine residues were hypothesized to exert a moderate effect (Fig. 4b; Table S7). Conserved FH-regulated cysteine residues were found to lie within functional domains of:

**Fig. 4.**
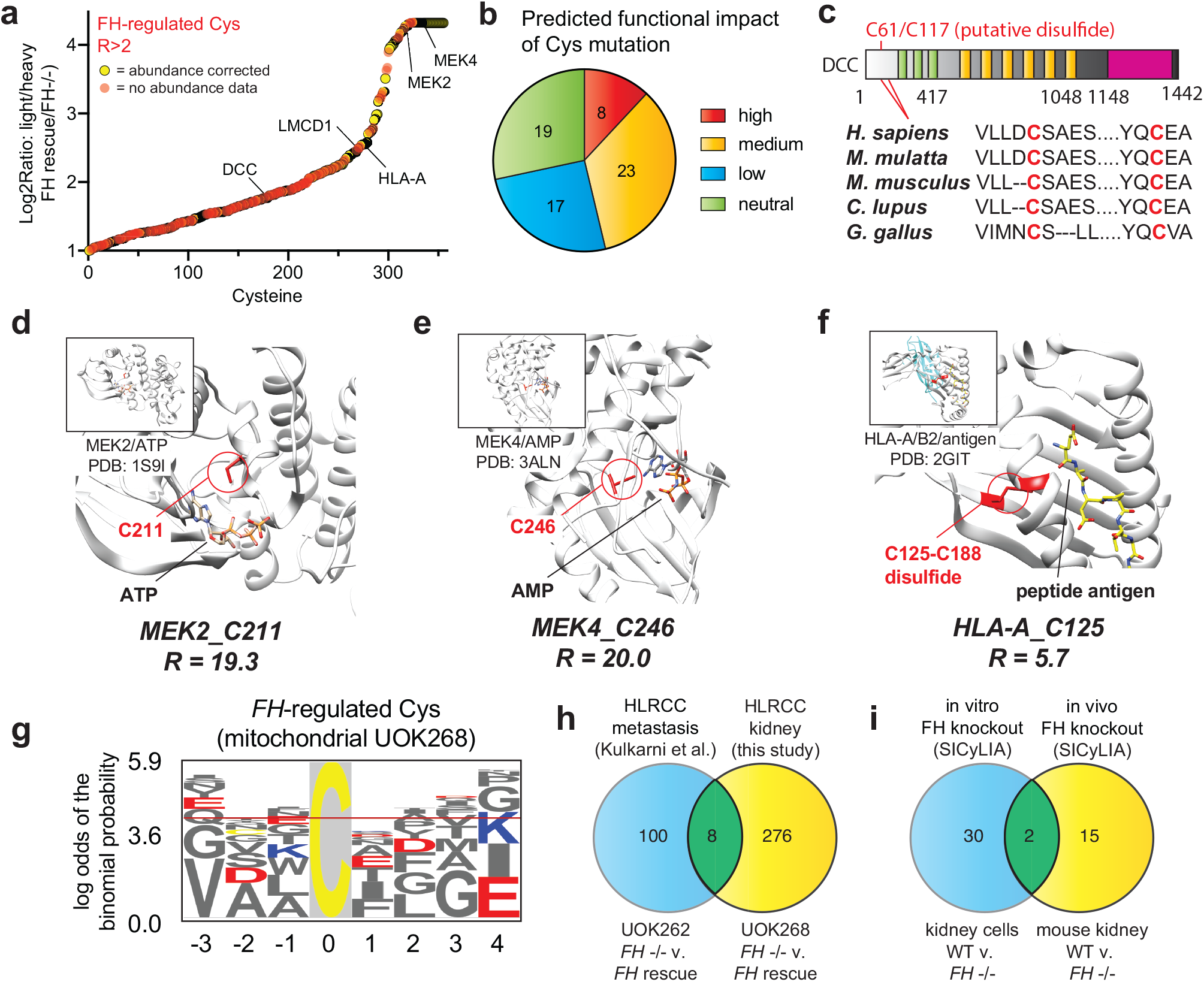
(a) Cysteines showing increased occupancy in FH rescue cells. Red = “High confidence” abundance corrected cysteine reactivity changes, Yellow = non-abundance corrected cysteine reactivity changes that potentially arise from either altered transcription or posttranslational modification. (b) Predicted functional impact of R>2 cysteines mapping to annotated cancer genes based on Mutation Assessor analysis. (c) Domain map of DCC, indicating conservation of C61 in metazoans. (d) Crystal structure of MEK2, showing position of FH-regulated cysteine near ATP-binding pocket. (e) Crystal structure of MEK4, showing position of FH-regulated cysteine near ATP-binding pocket. (f) Crystal structure of HLA-A, showing position of FH-regulated cysteine involved in a disulfide bond. (g) Motif analysis of FH-regulated cysteines. Logo is derived from mitochondrial cysteines, where fumarate concentrations are expected to be highest. Logos from cytosolic and nuclear FH-regulated cysteines are provided in Supplementary Information Fig. S2. (h) Overlap of FH-regulated cysteine residues identified in UOK262 and UOK268 cells. (i) Overlap of FH-regulated cysteine residues identified in mouse kidney cells and in vivo mouse kidney by van der Reest et al.

- **Tumor suppressors:** Examples include Deleted in Colon Cancer (DCC), in which FH-regulated Cys61 (C61, R=3.4), lies within a highly conserved region predicted to form a disulfide with Cys117 (Fig. 4c), and LIM and Cysteine Rich Domain-Containing Protein 1 Cys 52 (LCRDCP, C52, R=5.7), which is found in the Cys-rich region of the protein that is required for interaction and repression of GATA transcription factors.^30^
- **Signaling enzymes:** FH-regulated cysteines were identified within the ATP-binding sites of the MAP kinases MEK2 (C211, R=19.3, Fig. 4d) and MEK4 (C246, R=20, Fig. 4e), as well as within the catalytic subunit of AMPK (PRKAA2 C174, R=15.7 Table S6).^31^ The latter cysteine maps to a previously characterized site of functional oxidative modification in AMPK, suggesting a novel mechanistic input that may further reduce the AMPK activity of HLRCC tumors.^19^
- **Components of the immune system**: FH-dependent decreases in cysteine reactivity were observed in two Major Histocompatibility Complex (MHC) proteins: HLA-A (C125, R=3.0) and HLA-B (C188, R=3.8). Each map to an extracellular disulfide critical to the folding and structure of their respective MHC molecules (Fig. 4f),^32^ and the differential reactivity in the two cell lines indicates the potential of these proteins to be transcriptionally or posttranslationally regulated by FH.

Motif analysis of cysteine-containing peptides whose reactivity was decreased by ≥2-fold by *FH* inactivation found a high occurrence of flanking glutamate (E) and aspartate (D) residues, independent of protein localization (Fig. 4g, Fig. S2, Table S10). These carboxylate amino acids are not commonly found proximal to hyperreactive cysteines, and may contribute to fumarate reactivity via hydrogen bonding or stabilization of surface-exposed alpha helices.^17^ Overall, our survey of cysteine reactivity provides a novel resource for the identification of functional protein activity changes involved the development, progression, or treatment of HLRCC.

Concurrent with our initial chemoproteomic survey of *FH*-regulated cysteines in patient-derived UOK262 cells, Gottlieb et al. reported a related method for quantifying cysteine reactivity termed Stable Isotope Cysteine Labeling with IodoAcetamide (SICyLIA) and applied it to map cysteine occupancy changes in *Fh1* (termed *FH* here for brevity) knockout mice.^33^ SICyLIA differs from our approach in that it does not use an enrichment step to increase the coverage of low abundance cysteine residues, but similarly provides a quantitative and mechanism agnostic readout of *FH*-regulated cysteine reactivity. To understand how covalent S-succination and oxidative cysteine modifications vary across diverse models of *FH* inactivation, we examined overlap between four complementary datasets: UOK262 (*FH−/−* and FH rescue), UOK268 (*FH−/−* and FH rescue), mouse kidney cells (wild-type and *FH−/−*), and in vivo mouse kidney (wild-type and *FH−/−*).^17, 33^ Starting with analysis of *FH*-regulated cysteines in human UOK262/UOK268 cells, we found only eight sites with R≥2 overlap (Table S8, Fig. 4h). Included amongst these hits was Chromobox 5 (CBX5 C188, R=4.8), whose cysteine-dependent capture we have previously shown is sensitive to fumarate. Intrigued by the limited overlap in these HLRCC models, we performed a similar analysis of peptides found to exhibit 2-fold decreased reactivity (log2fold change >1) upon FH knockout in mouse kidney cells and mouse kidney tissues by SICyLIA (Table S9). Here again, *FH*-regulated cysteines between these two models demonstrated a very modest overlap, with only two specific cysteine residues (Cathepsin D C288 and Thioredoxin domain-containing protein 5, C293) displaying coordinate regulation by FH (Fig. 4i). No common residues were found to be regulated amongst all four models. Overall, these data demonstrate that considerable heterogeneity may be observed in the adaptive response of reactive cysteines to fumarate accumulation in different HLRCC models.

## Discussion

Here we have reported a quantitative chemoproteomic analysis of cysteine reactivity in the kidney-derived HLRCC UOK268 tumor cell line. Application of this approach enabled the identification of new candidate reactive cysteine residues potentially subject to posttranslational regulation by FH activity. Integrating this approach with quantitative whole proteome analyses of kidney-derived *FH−/−* and *FH+/+* rescue cells identified additional pathways upregulated at the level of protein expression in UOK268, including those related to the generation of mitochondrial NADH by the TCA cycle. An additional notable target identified in these studies were two variants of the MHC complex (HLA-A and HLA-B), which were found to undergo specific losses of reactivity in their extracellular regions. Previous studies have found the redox modification of HLA cysteines can result in altered antigen presentation.^32^ Our studies raise the possibility that reducing fumarate levels by *FH* rescue may augment the immune response, while *FH* inactivation may limit it. Together, these data provide a fertile mechanistic resource from which to formulate biological hypotheses regarding the development and progression of this genetically-defined kidney cancer.

An unexpected finding of our study was that proteomes derived from distinct physiological backgrounds demonstrate substantial heterogeneity in the adaptive response of reactive cysteine residues to fumarate accumulation. This is typified by the distinct cohort of downregulated reactive cysteines in UOK262 and UOK268 cells,^17^ as well as the limited overlap between *in vitro* and *in vivo* mouse models of FH loss studied by SICyLIA.^33^ This potentially reflects differences in the underlying biology of the tissues of origin, as well as the magnitude and spatial and temporal generation of fumarate in these different models. For example, unique cellular and subcellular levels of fumarate in these models would be expected to lead to distinct effects on S-succination and oxidative cysteine modifications, as well as transcriptional signaling in the nucleus. Comparative metabolomic analyses of these different models, and the development of improved tools to assess subcellular metabolite levels should aid in the future definition of this phenomenon.^34–35^ An ancillary insight from our analysis is the distinct information available from studying the consequences of *FH* rescue (as was done in this study) versus studying the conditions of *FH*-dependent transformation. For example, our whole proteome analyses unexpectedly found glycolytic enzymes become even more highly expressed in UOK268 cells upon FH rescue, suggesting that, at least in this model, the Warburg effect may not be completely reversible by re-introduction of FH alone. In the future, we anticipate that combining chemoproteomic profiling with methods to modulate FH levels with high temporal precision^36–37^ will greatly aid the identification of specific oncometabolite-driven cysteine reactivity changes which contribute to tumorigenesis.

Several recent studies have demonstrated that proteomic profiling of oncogene-induced tumorigenesis can aid the design of new treatment strategies,^38–40^ and two therapeutic hypotheses are raised by our data. The first is that targeting pathways which demonstrate a compensatory upregulation upon FH inactivation (either at the level of protein abundance or cysteine reactivity) may serve to limit adaptation of cancer cells and trigger cell death. The second is that inhibiting pathways found to be “damaged” by S-succination (R≥2) or downregulated upon FH loss at the whole proteome level may push essential pathways already poised on the precipice of failure over the edge, inducing cytotoxicity.^41^ Future studies integrating proteomics data with screens of pathway-targeted inhibitors^42^ and polypharmacological cysteine reactive fragments^43–44^ should enable the rapid testing of these hypotheses and facilitate the identification of new FH-dependent biology, therapeutic targets, and pre-clinical drug candidates. Overall, these studies extend our quantitative knowledge of the *FH*-dependent proteome and provide a novel resource to aid therapeutic design in HLRCC.

### Experimental Procedures

Full experimental procedures including cell culture, preparation of proteomes, LC-MS/MS, data analysis and Tables S1-S12 are provided in the Supplementary Information.

## Data accessibility

Proteomics data has been uploaded to the PRIDE database and can be accessed under identifier PDX009378.

## Acknowledgments

This work was supported by the Intramural Research Program of the NIH, National Cancer Institute, Center for Cancer Research (ZIA BC011488 and ZIA BC011038) and the CCR FLEX Program. Support for E.W. was provided by the NIH (1R01GM117004 and 1R01GM118431– 01A1).

## References

1. Liu, J. Y.; Wellen, K. E., Advances into understanding metabolites as signaling molecules in cancer progression. Curr Opin Cell Biol 2020, 63, 144–153.

2. Meier, J. L., Metabolic mechanisms of epigenetic regulation. ACS Chem Biol 2013, 8 (12), 2607–21.

3. Schvartzman, J. M.; Thompson, C. B.; Finley, L. W. S., Metabolic regulation of chromatin modifications and gene expression. J Cell Biol 2018, 217 (7), 2247–2259.

4. Kulkarni, R. A.; Montgomery, D. C.; Meier, J. L., Epigenetic regulation by endogenous metabolite pharmacology. Curr Opin Chem Biol 2019, 51, 30–39.

5. Harmel, R.; Fiedler, D., Features and regulation of non-enzymatic post-translational modifications. Nat Chem Biol 2018, 14 (3), 244–252.

6. Alderson, N. L.; Wang, Y.; Blatnik, M.; Frizzell, N.; Walla, M. D.; Lyons, T. J.; Alt, N.; Carson, J. A.; Nagai, R.; Thorpe, S. R.; Baynes, J. W., S-(2-Succinyl)cysteine: a novel chemical modification of tissue proteins by a Krebs cycle intermediate. Arch Biochem Biophys 2006, 450 (1), 1–8.

7. Bardella, C.; El-Bahrawy, M.; Frizzell, N.; Adam, J.; Ternette, N.; Hatipoglu, E.; Howarth, K.; O’Flaherty, L.; Roberts, I.; Turner, G.; Taylor, J.; Giaslakiotis, K.; Macaulay, V. M.; Harris, A. L.; Chandra, A.; Lehtonen, H. J.; Launonen, V.; Aaltonen, L. A.; Pugh, C. W.; Mihai, R.; Trudgian, D.; Kessler, B.; Baynes, J. W.; Ratcliffe, P. J.; Tomlinson, I. P.; Pollard, P. J., Aberrant succination of proteins in fumarate hydratase-deficient mice and HLRCC patients is a robust biomarker of mutation status. J Pathol 2011, 225 (1), 4–11.

8. Merkley, E. D.; Metz, T. O.; Smith, R. D.; Baynes, J. W.; Frizzell, N., The succinated proteome. Mass Spectrom Rev 2014, 33 (2), 98–109.

9. Sullivan, L. B.; Martinez-Garcia, E.; Nguyen, H.; Mullen, A. R.; Dufour, E.; Sudarshan, S.; Licht, J. D.; Deberardinis, R. J.; Chandel, N. S., The proto-oncometabolite fumarate binds glutathione to amplify ROS-dependent signaling. Mol Cell 2013, 51 (2), 236–48.

10. Zheng, L.; Cardaci, S.; Jerby, L.; MacKenzie, E. D.; Sciacovelli, M.; Johnson, T. I.; Gaude, E.; King, A.; Leach, J. D.; Edrada-Ebel, R.; Hedley, A.; Morrice, N. A.; Kalna, G.; Blyth, K.; Ruppin, E.; Frezza, C.; Gottlieb, E., Fumarate induces redox-dependent senescence by modifying glutathione metabolism. Nat Commun 2015, 6, 6001.

11. Paulsen, C. E.; Carroll, K. S., Cysteine-mediated redox signaling: chemistry, biology, and tools for discovery. Chem Rev 2013, 113 (7), 4633–79.

12. MacKenzie, E. D.; Selak, M. A.; Tennant, D. A.; Payne, L. J.; Crosby, S.; Frederiksen, C. M.; Watson, D. G.; Gottlieb, E., Cell-permeating alpha-ketoglutarate derivatives alleviate pseudohypoxia in succinate dehydrogenase-deficient cells. Mol Cell Biol 2007, 27 (9), 3282–9.

13. Tennant, D. A.; Frezza, C.; MacKenzie, E. D.; Nguyen, Q. D.; Zheng, L.; Selak, M. A.; Roberts, D. L.; Dive, C.; Watson, D. G.; Aboagye, E. O.; Gottlieb, E., Reactivating HIF prolyl hydroxylases under hypoxia results in metabolic catastrophe and cell death. Oncogene 2009, 28 (45), 4009–21.

14. Kinch, L.; Grishin, N. V.; Brugarolas, J., Succination of Keap1 and activation of Nrf2-dependent antioxidant pathways in FH-deficient papillary renal cell carcinoma type 2. Cancer Cell 2011, 20 (4), 418–20.

15. Adam, J.; Hatipoglu, E.; O’Flaherty, L.; Ternette, N.; Sahgal, N.; Lockstone, H.; Baban, D.; Nye, E.; Stamp, G. W.; Wolhuter, K.; Stevens, M.; Fischer, R.; Carmeliet, P.; Maxwell, P. H.; Pugh, C. W.; Frizzell, N.; Soga, T.; Kessler, B. M.; El-Bahrawy, M.; Ratcliffe, P. J.; Pollard, P. J., Renal cyst formation in Fh1-deficient mice is independent of the Hif/Phd pathway: roles for fumarate in KEAP1 succination and Nrf2 signaling. Cancer Cell 2011, 20 (4), 524–37.

16. Sciacovelli, M.; Goncalves, E.; Johnson, T. I.; Zecchini, V. R.; da Costa, A. S.; Gaude, E.; Drubbel, A. V.; Theobald, S. J.; Abbo, S. R.; Tran, M. G.; Rajeeve, V.; Cardaci, S.; Foster, S.; Yun, H.; Cutillas, P.; Warren, A.; Gnanapragasam, V.; Gottlieb, E.; Franze, K.; Huntly, B.; Maher, E. R.; Maxwell, P. H.; Saez-Rodriguez, J.; Frezza, C., Fumarate is an epigenetic modifier that elicits epithelial-to-mesenchymal transition. Nature 2016, 537 (7621), 544–547.

17. Kulkarni, R. A.; Bak, D. W.; Wei, D.; Bergholtz, S. E.; Briney, C. A.; Shrimp, J. H.; Alpsoy, A.; Thorpe, A. L.; Bavari, A. E.; Crooks, D. R.; Levy, M.; Florens, L.; Washburn, M. P.; Frizzell, N.; Dykhuizen, E. C.; Weerapana, E.; Linehan, W. M.; Meier, J. L., A chemoproteomic portrait of the oncometabolite fumarate. Nat Chem Biol 2019, 15 (4), 391–400.

18. Weerapana, E.; Wang, C.; Simon, G. M.; Richter, F.; Khare, S.; Dillon, M. B.; Bachovchin, D. A.; Mowen, K.; Baker, D.; Cravatt, B. F., Quantitative reactivity profiling predicts functional cysteines in proteomes. Nature 2010, 468 (7325), 790–5.

19. Tong, W. H.; Sourbier, C.; Kovtunovych, G.; Jeong, S. Y.; Vira, M.; Ghosh, M.; Romero, V. V.; Sougrat, R.; Vaulont, S.; Viollet, B.; Kim, Y. S.; Lee, S.; Trepel, J.; Srinivasan, R.; Bratslavsky, G.; Yang, Y.; Linehan, W. M.; Rouault, T. A., The glycolytic shift in fumarate-hydratase-deficient kidney cancer lowers AMPK levels, increases anabolic propensities and lowers cellular iron levels. Cancer Cell 2011, 20 (3), 315–27.

20. Yang, Y.; Valera, V.; Sourbier, C.; Vocke, C. D.; Wei, M.; Pike, L.; Huang, Y.; Merino, M. A.; Bratslavsky, G.; Wu, M.; Ricketts, C. J.; Linehan, W. M., A novel fumarate hydratase-deficient HLRCC kidney cancer cell line, UOK268: a model of the Warburg effect in cancer. Cancer Genet 2012, 205 (7-8), 377–90.

21. Hsu, J. L.; Huang, S. Y.; Chow, N. H.; Chen, S. H., Stable-isotope dimethyl labeling for quantitative proteomics. Anal Chem 2003, 75 (24), 6843–52.

22. Bar-Peled, L.; Kemper, E. K.; Suciu, R. M.; Vinogradova, E. V.; Backus, K. M.; Horning, B. D.; Paul, T. A.; Ichu, T. A.; Svensson, R. U.; Olucha, J.; Chang, M. W.; Kok, B. P.; Zhu, Z.; Ihle, N. T.; Dix, M. M.; Jiang, P.; Hayward, M. M.; Saez, E.; Shaw, R. J.; Cravatt, B. F., Chemical Proteomics Identifies Druggable Vulnerabilities in a Genetically Defined Cancer. Cell 2017, 171 (3), 696–709 e23.

23. Huang da, W.; Sherman, B. T.; Lempicki, R. A., Systematic and integrative analysis of large gene lists using DAVID bioinformatics resources. Nat Protoc 2009, 4 (1), 44–57.

24. Grubb, R. L., 3rd; Franks, M. E.; Toro, J.; Middelton, L.; Choyke, L.; Fowler, S.; Torres-Cabala, C.; Glenn, G. M.; Choyke, P.; Merino, M. J.; Zbar, B.; Pinto, P. A.; Srinivasan, R.; Coleman, J. A.; Linehan, W. M., Hereditary leiomyomatosis and renal cell cancer: a syndrome associated with an aggressive form of inherited renal cancer. J Urol 2007, 177 (6), 2074–9; discussion 2079-80.

25. Xu, Y.; Andrade, J.; Ueberheide, B.; Neel, B. G., Activated Thiol Sepharose-based proteomic approach to quantify reversible protein oxidation. FASEB J 2019, 33 (11), 12336–12347.

26. Reva, B.; Antipin, Y.; Sander, C., Predicting the functional impact of protein mutations: application to cancer genomics. Nucleic Acids Res 2011, 39 (17), e118.

27. Repana, D.; Nulsen, J.; Dressler, L.; Bortolomeazzi, M.; Venkata, S. K.; Tourna, A.; Yakovleva, A.; Palmieri, T.; Ciccarelli, F. D., The Network of Cancer Genes (NCG): a comprehensive catalogue of known and candidate cancer genes from cancer sequencing screens. Genome Biol 2019, 20 (1), 1.

28. Gao, J.; Aksoy, B. A.; Dogrusoz, U.; Dresdner, G.; Gross, B.; Sumer, S. O.; Sun, Y.; Jacobsen, A.; Sinha, R.; Larsson, E.; Cerami, E.; Sander, C.; Schultz, N., Integrative analysis of complex cancer genomics and clinical profiles using the cBioPortal. Sci Signal 2013, 6 (269), pl1.

29. Zhang, J.; Manley, J. L., Misregulation of pre-mRNA alternative splicing in cancer. Cancer Discov 2013, 3 (11), 1228–37.

30. Rath, N.; Wang, Z.; Lu, M. M.; Morrisey, E. E., LMCD1/Dyxin is a novel transcriptional cofactor that restricts GATA6 function by inhibiting DNA binding. Mol Cell Biol 2005, 25 (20), 8864–73.

31. Shao, D.; Oka, S.; Liu, T.; Zhai, P.; Ago, T.; Sciarretta, S.; Li, H.; Sadoshima, J., A redox-dependent mechanism for regulation of AMPK activation by Thioredoxin1 during energy starvation. Cell Metab 2014, 19 (2), 232–45.

32. Trujillo, J. A.; Croft, N. P.; Dudek, N. L.; Channappanavar, R.; Theodossis, A.; Webb, A. I.; Dunstone, M. A.; Illing, P. T.; Butler, N. S.; Fett, C.; Tscharke, D. C.; Rossjohn, J.; Perlman, S.; Purcell, A. W., The cellular redox environment alters antigen presentation. J Biol Chem 2014, 289 (40), 27979–91.

33. van der Reest, J.; Lilla, S.; Zheng, L.; Zanivan, S.; Gottlieb, E., Proteome-wide analysis of cysteine oxidation reveals metabolic sensitivity to redox stress. Nat Commun 2018, 9 (1), 1581.

34. Wellen, K. E.; Snyder, N. W., Should we consider subcellular compartmentalization of metabolites, and if so, how do we measure them? Curr Opin Clin Nutr Metab Care 2019, 22 (5), 347–354.

35. Kulkarni, R. A.; Briney, C. A.; Crooks, D. R.; Bergholtz, S. E.; Mushti, C.; Lockett, S. J.; Lane, A. N.; Fan, T. W.; Swenson, R. E.; Marston Linehan, W.; Meier, J. L., Photoinducible Oncometabolite Detection. Chembiochem 2019, 20 (3), 360–365.

36. Nabet, B.; Roberts, J. M.; Buckley, D. L.; Paulk, J.; Dastjerdi, S.; Yang, A.; Leggett, A. L.; Erb, M. A.; Lawlor, M. A.; Souza, A.; Scott, T. G.; Vittori, S.; Perry, J. A.; Qi, J.; Winter, G. E.; Wong, K. K.; Gray, N. S.; Bradner, J. E., The dTAG system for immediate and target-specific protein degradation. Nat Chem Biol 2018, 14 (5), 431–441.

37. Natsume, T.; Kiyomitsu, T.; Saga, Y.; Kanemaki, M. T., Rapid Protein Depletion in Human Cells by Auxin-Inducible Degron Tagging with Short Homology Donors. Cell Rep 2016, 15 (1), 210–218.

38. Santana-Codina, N.; Chandhoke, A. S.; Yu, Q.; Malachowska, B.; Kuljanin, M.; Gikandi, A.; Stanczak, M.; Gableske, S.; Jedrychowski, M. P.; Scott, D. A.; Aguirre, A. J.; Fendler, W.; Gray, N. S.; Mancias, J. D., Defining and Targeting Adaptations to Oncogenic KRAS(G12C) Inhibition Using Quantitative Temporal Proteomics. Cell Rep 2020, 30 (13), 4584–4599 e4.

39. Biancur, D. E.; Paulo, J. A.; Malachowska, B.; Quiles Del Rey, M.; Sousa, C. M.; Wang, X.; Sohn, A. S. W.; Chu, G. C.; Gygi, S. P.; Harper, J. W.; Fendler, W.; Mancias, J. D.; Kimmelman, A. C., Compensatory metabolic networks in pancreatic cancers upon perturbation of glutamine metabolism. Nat Commun 2017, 8, 15965.

40. Kohnz, R. A.; Mulvihill, M. M.; Chang, J. W.; Hsu, K. L.; Sorrentino, A.; Cravatt, B. F.; Bandyopadhyay, S.; Goga, A.; Nomura, D. K., Activity-Based Protein Profiling of Oncogene-Driven Changes in Metabolism Reveals Broad Dysregulation of PAFAH1B2 and 1B3 in Cancer. ACS Chem Biol 2015, 10 (7), 1624–30.

41. Muller, F. L.; Aquilanti, E. A.; DePinho, R. A., Collateral Lethality: A new therapeutic strategy in oncology. Trends Cancer 2015, 1 (3), 161–173.

42. Yohe, M. E.; Gryder, B. E.; Shern, J. F.; Song, Y. K.; Chou, H. C.; Sindiri, S.; Mendoza, A.; Patidar, R.; Zhang, X.; Guha, R.; Butcher, D.; Isanogle, K. A.; Robinson, C. M.; Luo, X.; Chen, J. Q.; Walton, A.; Awasthi, P.; Edmondson, E. F.; Difilippantonio, S.; Wei, J. S.; Zhao, K.; Ferrer, M.; Thomas, C. J.; Khan, J., MEK inhibition induces MYOG and remodels super-enhancers in RAS-driven rhabdomyosarcoma. Sci Transl Med 2018, 10 (448).

43. Backus, K. M.; Correia, B. E.; Lum, K. M.; Forli, S.; Horning, B. D.; Gonzalez-Paez, G. E.; Chatterjee, S.; Lanning, B. R.; Teijaro, J. R.; Olson, A. J.; Wolan, D. W.; Cravatt, B. F., Proteome-wide covalent ligand discovery in native biological systems. Nature 2016, 534 (7608), 570–4.

44. Counihan, J. L.; Wiggenhorn, A. L.; Anderson, K. E.; Nomura, D. K., Chemoproteomics-Enabled Covalent Ligand Screening Reveals ALDH3A1 as a Lung Cancer Therapy Target. ACS Chem Biol 2018, 13 (8), 1970–1977.

