## Supplementary Information for "Heterogeneous adaptation of cysteine reactivity to a covalent oncometabolite"

Heterogeneous Adaptation of Cysteine Reactivity to a Cov­alent Oncometabolite

**Table of Contents for Supporting Information**

Table of Contents	S1

Supplementary Figures S1-S2	S2-3

Cell culture and isolation of whole cell lysates	S4

Chemoproteomic analysis of *FH*-regulated cysteines	S5-7

Whole proteome protein ReDiMe abundance analysis	S8-9

Bioinformatic analysis of FH-regulated cysteines	S10

References	S11

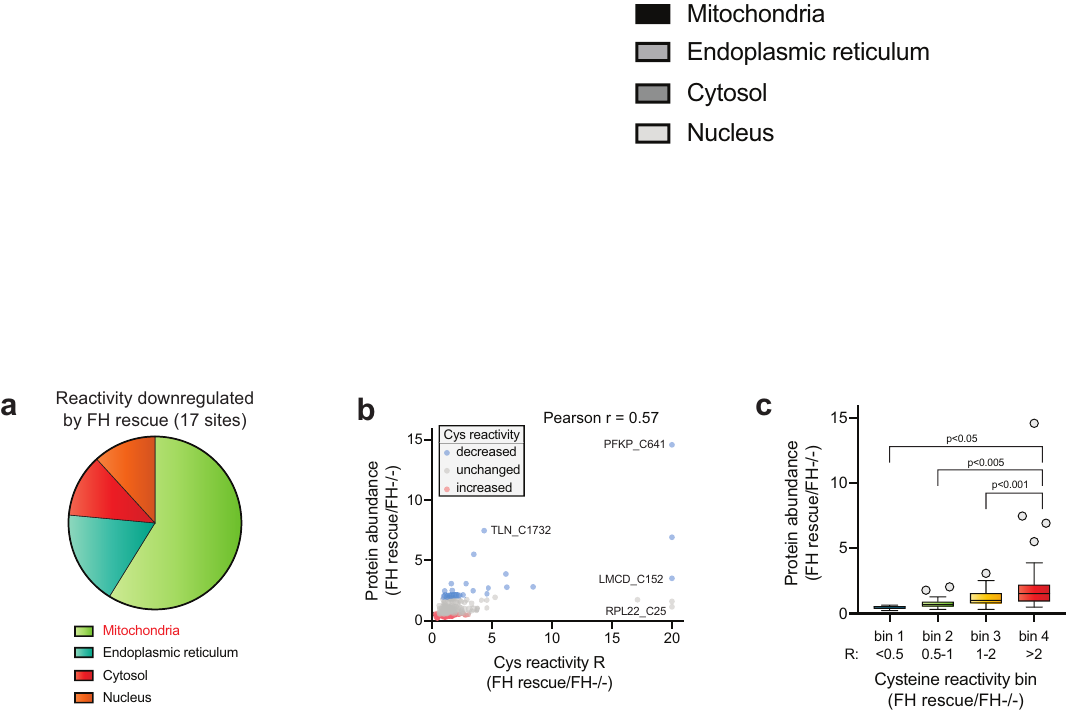

**Figure S1** (a) Cellular localization of cysteine residues downregulatedd by FH rescue (R≥2, n≥2 SD≤50%). (b) Correlation of protein abundance and cysteine reactivity (n≥2 SD≤25%). (c) Cysteines with greater reactivity in FH rescue cells (right bin, red) have significantly increased protein abundance relative to all other cysteine reactivity subsets (R≥2, n≥2 SD≤25%).

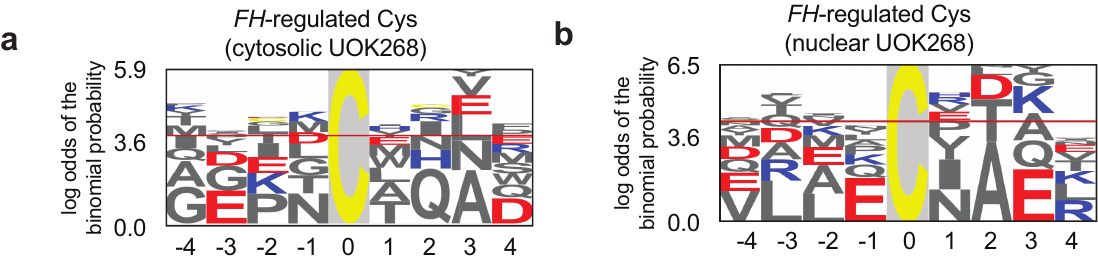

**Figure S2** Motif analysis of FH-regulated cysteines derived from (a) cytosolic, and (b) nuclear proteins.

**Cell culture and isolation of whole cell lysates**

UOK268 (FH -/-) cells were cultured at 37 °C under 5% CO_2_ atmosphere grown in Dulbecco’s modified eagle medium (DMEM) supplemented with 10 % fetal bovine serum, 2 mM L-glutamine and 1 mM sodium pyruvate, while the UOK268 FH (+/+) rescue media was supplemented with 0.3 µg/mL Blasticidin. Cells were grown to 80% confluency then DMEM media was removed and cells were washed with PBS. Proteomes were harvested by rinsing cells 2X with ice cold PBS and scrapped into a 50 mL conical. Subsequently, cells were washed 3X with 30 mL of ice-cold PBS and centrifuged (500 r.c.f x 5 min, 4°C). Following the final PBS wash, cell pellets were resuspended with 1 mL of ice-cold PBS, transferred to a 1.7 mL microtube and centrifuged to remove excess PBS. Cell pellets were resuspended with 1 mL of lysis buffer consisting of PBS with 1X protease inhibitor cocktail (Cell Signaling # 5871). The resuspended cells were lysed by sonication (3 x 1 second pulse, amplitude 1, 60 seconds resting on ice between pulses) using a 100W QSonica XL2000 sonicator. Following sonication, the lysates were centrifuged (14,000 r.c.f x 30 minutes, 4 °C) and protein concentration was determined via Qubit Protein Assay Kit on a Qubit 2.0 Fluorometer. Lysates were diluted with lysis buffer to a 2 mg/mL protein concentration and store at -80 °C.

**Chemoproteomic analysis of *FH*-regulated cysteines**

*Chemoproteomic labeling and enrichment of FH-regulated cysteines*

For identification and enrichment of *FH*-regulated cysteines (Table S1), 2 mg of UOK268 (FH-/-) or UOK268WT (FH+/+) proteomes (1 mL, 2 mg/mL) were labeled with 100 µM IA-alkyne (10 μL, 10 mM stock in DMSO) for 1 h at room temperature. Probe labeled proteins were then conjugated to UV-cleavable biotin-azide (UV-azo) tag by Cu(I)-catalyzed [3 + 2] cycloaddition as previously reported.^1^ Briefly, azo-tag (100 μM), TCEP (1 mM), TBTA (100 μM), and CuSO_4_ (1 mM) were sequentially added to the labeled proteome. Reactions were vortexed and incubated at room temperature for 1 h. Proteomes were centrifuged (6500 rcf *x* 10 min, 4 °C) to collect precipitated protein. Supernatant was discarded, and protein pellets were resuspended in 500 μL of methanol (dry-ice chilled) with sonication, and centrifuged (6500 rcf *x* 10 min, 4 °C). This step was repeated, and the resulting washed pellets was redissolved (1.2% w/v SDS in PBS; 1 mL); sonication followed by heating at 80-95 °C for 5 min was used to ensure complete solubilization. Samples were cooled to room temperature, diluted with PBS (5.0 mL), and incubated with Streptavidin beads (100 μL of 50% aqueous slurry per enrichment) overnight at 4 °C. Samples were allowed to warm to room temperature, pelleted by centrifugation (1400 rcf *x* 3 min), and supernatant discarded. Beads were then sequentially washed with 0.2% SDS in PBS (5 mL x 1), PBS (5 mL x 3) and H_2_O (5 mL x 3) for a total of 7 washes.

*On-bead reductive alkylation, and tryptic digest of proteomic samples*

Following the final wash, protein-bound streptavidin beads were resuspended in 6 M urea in PBS (500 μL) and reductively alkylated by sequential addition of 10 mM DTT (25 μL of 200 mM in H_2_O, 65 °C for 20 min) and 20 mM iodoacetamide (25 μL of 400 mM in H_2_O, 37 °C for 30 min) to each sample. Reactions were then diluted by addition of PBS (950 μL), pelleted by centrifugation (1400 rcf x 3 min), and the supernatant discarded. Samples were then subjected to tryptic digest by addition of 200 μL of a pre-mixed solution of 2M urea in PBS, 1 mM CaCl_2_ (2 μL of 100 mM in H_2_O), and 2 μg of Trypsin Gold (Promega, 4 μL of 0.5 μg/μL in 1% acetic acid). Samples were shaken overnight at 37 °C and pelleted by centrifugation (1400 rcf x 3 min). Beads were then washed sequentially with PBS (500 µL x 2), H_2_O (500 µL x 2), and 100 mM triethylammonium bicarbonate (TEAB) buffer (500 µL x 2) and subsequently resuspended in 100 mM TEAB (100 µL).

*On-bead reductive dimethylation (ReDiMe) and UV-cleavage of proteomic samples*

Isotopic labeling of samples was achieved through on-bead reductive dimethylation (ReDiMe),^2^ performed by the addition of 4 µL of light (UOK268, FH+/+) or heavy (UOK268, FH-/-) 20% formaldehyde (HCHO or D^13^CDO, respectively) and 20 µL of 0.6 M sodium cyanoborohydride (25 ºC, 2 hours). Beads were then washed with 100 mM TEAB (500 µL x 2), mixed, and washed with H_2_O (500 µL x 2). The combined beads were resuspended in H_2_O (200 µL) and irradiated with 365 nm UV light for 1 h to release labeled peptides. Beads were pelleted by centrifugation (1,400 g, 3 min, 25 ºC) and the supernatant transferred to a new centrifuge tube. The beads were washed with PBS (100 µL, 2x), with the washes being combined with the previous supernatants, for a total eluted peptide sample volume of ~ 350 µL. Formic acid (17.5 µL) was added to a final concentration of 5% and samples were stored at -20 ºC until ready for LC/LC-MS/MS analysis.

*LC/LC-MS/MS and data analysis for quantitative cysteine reactivity profiling*

Mass spectrometry was performed using a Thermo LTQ Orbitrap Discovery mass spectrometer coupled to an Agilent 1200 series HPLC. Labeled peptide samples were pressure loaded onto 250 mm fused silica desalting column packed with 4 cm of Aqua C18 reverse phase resin (Phenomenex). Peptides were eluted onto a 100-mm fused silica biphasic column packed with 10 cm C18 resin and 4 cm Partisphere strong cation exchange resin (SCX, Whatman), using a five-step multidimensional LC-MS protocol (MudPIT). Each of the five steps used a salt push (0%, 50%, 80%, 100%, and 100%), followed by a gradient of buffer B in Buffer A (Buffer A: 95% water, 5% acetonitrile, 0.1% formic acid; Buffer B: 20% water, 80% acetonitrile, 0.1% formic acid) as outlined previously.^3^ The flow rate through the column was ~0.25 µL/min, with a spray voltage of 2.75 kV. One full MS1 scan (400-1800 MW) was followed by 8 data dependent scans of the n^th^ most intense ion. Dynamic exclusion was enabled. The tandem MS data, generated from the 5 MudPIT runs, was analyzed by the SEQUEST algorithm.^4^ Both static (+57.0215 m/z, iodoacetamide alkylation) and differential (+180.1375 m/z, iodacetamide alkyne labeling) modifications on cysteine residues were specified. The precursor-ion mass tolerance was set at 50 ppm while the fragment-ion mass tolerance was set to 0 (default setting). Data was searched against a human reverse-concatenated non-redundant FASTA database containing Uniprot identifiers. MS datasets were independently searched with light and heavy ReDiMe parameter files; for these searches, static modifications on lysine and the n-terminus of +28.0313 (light) or +34.0632 (heavy) were used. MS2 spectra matches were assembled into protein identifications and filtered using DTASelect2.0,^5^ to generate a list of protein hits with a peptide false-discovery rate of <5%. With the –trypstat and –modstat options applied, peptides were restricted to fully tryptic (-y 2) with a found modification (-m 0) and a delta-CN score greater than 0.06 (-d 0.06). Single peptides per locus were also allowed (-p 1) as were redundant peptides identifications from multiple proteins, but the database contained only a single consensus splice variant for each protein. Quantification of peptide L/H ratios were calculated using the cimage quantification package described previously.^3^

**Whole proteome protein ReDiMe abundance analysis**

*Whole-proteome ReDiMe sample preparation*

For quantification of protein abundance changes upon FH-rescue (Table S3), 100 µg of both UOK268, FH-/- and UOK268, FH+/+ proteomes (100 µL, 1 mg/mL) were precipitated by the addition of 5 µL 100% trichloroacetic acid in PBS, vortexed and frozen (-80 ºC, overnight). After thawing, proteins were pelleted by centrifugation (15,000 g, 10 min, 4 ºC) and solvent removed. Protein pellets were resuspended in 500 µL of ice-cold acetone by bath sonication and pelleted by centrifugation (5000g, 10 min, 4 ºC). The solvent was removed and the pellet was allowed to air dry and then resuspended in 8 M Urea in 100 mM TEAB (30 µL). Reductive alkylation was performed by the sequential addition of 100 mM TEAB (70 µL), 1.5 µL of 1 M DTT (65 ºC, 15 min), and 2.5 µL of 400 mM iodoacetamide (25 ºC, 30 min). Reactions were diluted with additional 100 mM TEAB (120 µL) and tryptic digestion performed by the addition of 2 µg of sequencing-grade trypsin (4 µL of 20 µg diluted in H_2_O) and 2.5 µL of 100 mM CaCl_2_ (37 ºC, overnight). After tryptic digest, reductive dimethylation^6^ was performed by the addition of 4 µL of 20% light (UOK268, FH+/+) or heavy (UOK268, FH-/-) formaldehyde and 20 µL of 0.6 M sodium cyanoborohydride (25 ºC, 2 hours). The reaction was quenched by the addition of 8µL ammonium hydroxide (25 ºC, 15 min). The light (UOK268, FH+/+) and heavy (UOK268, FH-/-) tryptic peptide samples were then combined, desalted on a Sep-Pak, and dried by speed-vac. Peptide samples were stored at -20 ºC until ready for fractionation and LC-MS/MS analysis.

*Off-line high pH ReDiMe peptide fractionation*

Samples were resuspended in 500 µL of high pH buffer A (95% H_2_O, 5% acetonitrile, 10 mM ammonium bicarbonate) and loaded onto a manual injection loop connected to an Agilent 1100 Series HPLC. Peptides were separated on a 25 cm Agilent Extend-C18 column using a 60 min gradient from 20-35% high pH buffer B (10% H2O, 90% acetonitrile, 10 mM ammonium bicarbonate).^7^ Fractions were collected using a Gilson FC203B fraction collector into a 96 deep-well plate (0.6 min/well). Subsequent concatenation of every sixth well resulted in six pooled fractions for later LC-MS/MS analysis. These six fractions were dried by speed-vac and then resuspended in 30 µL of low pH buffer A (95% H_2_O, 5% acetonitrile, 0.1% formic acid).

*LC-MS/MS and data analysis for ReDiMe whole-proteome quantification*

LC-MS/MS was performed on a Thermo LTQ Oribtrap XL mass spectrometer (Thermo Scientific) coupled to an easy-nanoLC system (Agilent). For each of the six off-line fractions, 10 µL of peptide mixture was pre-loaded onto a C18 pre-column. Peptides were eluted onto a 100-mm fused silica column packed with 10 cm C18 resin using a 4 hour gradient of buffer B in Buffer A (Buffer A: 95% water, 5% acetonitrile, 0.1% formic acid; Buffer B: 20% water, 80% acetonitrile, 0.1% formic acid). The flow rate through the column was ~0.4 µL/min, with a spray voltage of 2.75 kV. One full MS1 scan (400-1800 MW) was followed by 8 data dependent scans of the n^th^ most intense ion. Dynamic exclusion was enabled. The tandem MS data was analyzed by the SEQUEST algorithm.^4^ Static modification of cysteine residues (+57.0215 m/z, iodoacetamide alkylation) was assumed with no enzyme specificity. The precursor-ion mass tolerance was set at 50 ppm while the fragment-ion mass tolerance was set to 0 (default setting). Data was searched against a human reverse-concatenated non-redundant FASTA database containing Uniprot identifiers. MS datasets were independently searched with light and heavy ReDiMe parameter files; for these searches, static modifications on lysine and the n-terminus of +28.0313 (light) or +34.0632 (heavy) were used. MS2 spectra matches were assembled into protein identifications and filtered using DTASelect2.0,^5^ to generate a list of protein hits with a peptide false-discovery rate of <5%. Quantification of protein L/H ratios were calculated using the cimage quantification package described previously.^3^

**Bioinformatic analysis of FH-regulated cysteines**

Annotation of protein subcellular localization as well as cysteine function and conservation was generated from the Uniprot Protein Knowledgebase (UniProtKB) as described previously.^8^ Analysis of linear sequences flanking *FH*-regulated cysteines was performed using the informatics tool pLogo, accessible at: [https://plogo.uconn.edu](https://plogo.uconn.edu/). Input sequences are listed in Table S10, and were derived from the 25 cysteines found to be most *FH*-regulated in each compartment (highest R values, n≥2, SD≤50%) using high-confidence values derived from ReDiMe abundance correction (Table S4). Protein sequences for motif analysis were derived from their tryptic peptide sequences using Peptide Extender ([schwartzlab.uconn.edu/pepextend](http://schwartzlab.uconn.edu/pepextend)). Conservation and functional impact of *FH*-regulated cysteines identified in chemoproteomic experiments was analyzed using the informatics tool Mutation Assessor, accessible at: <http://mutationassessor.org/r3>. Potential functional impact of fumarate modifications reflects the effect of C to G mutations on the functional impact (FI) output of Mutation Assessor. Gene ontology analysis was performed using the bioinformatics tool DAVID, accessible at: <http://david.ncifcrf.gov/>.

**References**

1. Qian, Y., Martell, J., Pace, N.J., Ballard, T.E., Johnson, D.S. and Weerapana, E., 2013. An isotopically tagged azobenzene‐based cleavable linker for quantitative proteomics. *Chembiochem*, *14*(12), pp.1410-1414.
2. Yang, F., Gao, J., Che, J., Jia, G. and Wang, C., 2018. A dimethyl-labeling-based strategy for site-specifically quantitative chemical proteomics. *Analytical chemistry*, *90*(15), pp.9576-9582.
3. Weerapana, E. *et al.* Quantitative reactivity profiling predicts functional cysteines in proteomes. *Nature* **468**, 790-795, doi:10.1038/nature09472 (2010).
4. Eng, J. K., McCormack, A. L. & Yates, J. R. An approach to correlate tandem mass spectral data of peptides with amino acid sequences in a protein database. *J Am Soc Mass Spec* **5**, 976-989 (1994).
5. Tabb, D. L., McDonald, W. H. & Yates, J. R. DTASelect and Contrast: tools for assembling and comparing protein identifications from shotgun proteomics. *J Proteom Res* **1**, 21-26 (2002).
6. Boersema, P.J., Raijmakers, R., Lemeer, S., Mohammed, S. and Heck, A.J., 2009. Multiplex peptide stable isotope dimethyl labeling for quantitative proteomics. *Nature protocols*, *4*(4), p.484.
7. Edwards, A. and Haas, W., 2016. Multiplexed quantitative proteomics for high-throughput comprehensive proteome comparisons of human cell lines. In *Proteomics in Systems Biology* (pp. 1-13). Humana Press, New York, NY.
8. Bak, D. W., Pizzagalli M. D., and Weerapana E. Identifying Functional Cysteine Residues in the Mitochondria. *ACS Chem Biol* **12**, 947–957 (2017).
